# MicroRNA775 and target *Galactosyltransferase* (*GALT9*) module regulates recovery from submergence induced senescence by modulating *SAGs* in *Arabidopsis thaliana*

**DOI:** 10.1101/2021.01.29.428849

**Authors:** Vishnu Mishra, Archita Singh, Nidhi Gandhi, Shabari Sarkar Das, Sandeep Yadav, Ashutosh Kumar, Ananda K. Sarkar

## Abstract

Submergence induced hypoxic condition is one of the abiotic stresses which negatively affects the plant growth and development, and causes early onset of senescence. Hypoxic conditions ateres the expression of a number of non-coding microRNAs (miRNAs), besides protein-coding genes. However, the molecular function of stress-induced miRNA in submergence induced physiological or developmental changes and recovery remains to be understood. The expression of miR775 is highly induced under hypoxic stress conditions. Here, we show that miR775 is a potential post-transcriptional regulator number of targets, including *Galactosyltransferase* (*GALT9*). The expression of miR775 and target *GALT9* was significantly induced and reduced respectively at 24 hours of submergence. The overexpression of miR775 (miR775-Oe) confers enhanced recovery from submergence stress and reduced accumulation of ROS, in contrast to wild type and endogenous target mimic of miR775 (*MIM775) Arabidopsis* plants. We observed a similar recovery phenotype in case of target *galt9* mutant plants, indicating the role of miR775-*GALT9* module in recovery from submergence. Further, we showed that the expression of *SENESCENCE ASSOCIATED GENES* (*SAGs*), such as *SAG12, SAG29*, and *ORE1*. was increased in *MIM775* and reduced in miR775-Oe and *galt9* plants. Thus, our results suggest that miR775-*GALT9* module plays a crucial role in the recovery from submergence by modulating the expression of *SAGs* through differential accumulation of ROS.

## Introduction

Abiotic factors like light, temperature, water, oxygen, nutrition, etc. act as external cues to modulate plant growth and development. In the due course of time, plants have evolved several adaptive features like spine-like leaves, sunken stomata, wax deposition on leaf, chlorophyll deposition in stem etc. to successfully complete their life cycle under diverse environmental conditions (Kraehmer, 2016). Globally, the flood is one of the major stresses that lead to severe loss in crop yield and productivity (Ismail, 2018, Kumar and Dash, 2019, Bui *et al*., 2020). Flooding stress is categorized into two categories depending upon the water exposure such as, complete submergence, and partial submergence. Under complete submergence, the whole plant is fully immersed in water, while under partial submergence, the shoot terminal is maintained above the water surface (Fukao *et al*., 2019). Premature senescence, necrosis, and chlorosis, cessation of growth are the major consequences of submergence stress (Zhang *et al*., 2000, Visser *et al*., 2003). Submergence initiates various molecular cascades that adversely affect plant growth and development. Submergence also leads to excessive reactive oxygen species (ROS) production and cell death due to the unavailability of oxygen (hypoxic conditions). Prolonged submergence gradually affects gaseous exchange which leads to cell damage and chlorophyll breakdown causing early onset of senescence (Fukao *et al*., 2019). *RESPIRATORY BURST OXIDASE HOMOLOGUE D* (*RBOHD*), is one of the NAPDH oxidases involved in the production of ROS. and is widely used as a marker gene to access the ROS accumulation during submergence stress. *ROHBD* promotes alcohol dehydrogenase (ADH), pyruvate decarboxylase 1 (PDC), lactate dehydrogenase (LDH), Calcium (Ca^2+^) levels and various hypoxia-responsive genes, induced during hypoxia stress. Abundance of *RBOHD* also changes under submergence stress (Yamauchi *et al*., 2013, Yeung *et al*., 2018). Some other regulatory genes are combinedly involved in flooding and submergence through different regulatory pathways during stress for example *SENESCENCE ASSOCIATED GENE* (*SAGs*) *SAG12, SAG29, ORESARA1 (ORE1/NAC6). SAG12* encodes a cysteine protease in *Arabidopsis*. The *sag12* mutant shows a decrease in yield and nitrogen content compared to the wild type (Col-0) in nitrogen deficiency and majorly involved in senescence (James *et al*., 2018). *SAG12* is mainly expressed in root stele and at the reproductive stage, especially in nitrogen deficiency conditions (James *et al*., 2018). In *Arabidopsis*, the plasma membrane-localized *Medicago truncatula* (MtN3 protein), *SAG29* modulates cell viability under high salinity. The *SAG29* gene is prominently expressed in senescing plant tissues. The *SAG29* gene is highly induced by osmotic stresses through an abscisic acid-dependent pathway. The *SAG29* overexpressing transgenic line shows more senescence and is hypersensitive to salt stress compared to the *sag29* mutant (Seo *et al*., 2011b). *ORE1* has been identified as an accession specific regulatory gene which is highly induced in Bay-0 ecotype and has a predominant role in chlorophyll breakdown (Yeung *et al*., 2018). The knockout mutants of *ore1* in the Col-0 background show intermediary submergence tolerance (Yeung *et al*., 2018).

Besides genes and TFs, a large number of miRNAs have been also reported to be dynamically upregulated and downregulated under submergence stress (Zhang *et al*., 2008, Moldovan *et al*., 2010, Licausi *et al*., 2011, Liu *et al*., 2012, Jeong *et al*., 2013, Zhai *et al*., 2013, Jin *et al*., 2017, Li *et al*., 2017, Franke *et al*., 2018, Fukao *et al*., 2019). MicroRNAs belong to a class of endogenous small non-coding RNAs, which negatively regulate their target genes expression at the post-transcriptional level through complementary pairing with their specific target mRNAs in most of the eukaryotes. Several miRNAs have recently been implicated in various developmental processes, including shoot and root development. Few miRNAs have been shown to be differentially expressed under various abiotic stresses conditions (Singh *et al*., 2020a, Wang *et al*., 2020).

In *Arabidopsis*, the NAC transcription factor *ORE1* regulates approximately 46% of the 170 genes that are known to be senescence-associated genes indicating *ORE1* is a key regulatory gene involved in senescence. miR164 mediated regulation of *ORE1* triggers early senescence by controlling various *SAGs* gene (Balazadeh *et al*., 2010, Glazińska *et al*., 2014, Yeung *et al*., 2018). miR159 was upregulated in maize root during waterlogging and was responsible for flood regulation by modulating their target *GAMYBs, MYB33*, and *MYB101* homologs (Liu *et al*., 2012). miR166 is shown to have an important role in response to flood stress through regulating calcium spikes and accumulation of ROS during root growth and development (Fukao *et al*., 2019). miR167 was found to regulate short-term waterlogging or submergence in maize root by targeting *ARFs* (Zhang *et al*., 2008, Liu *et al*., 2012). Recently, it has been reported that miR167 was differentially upregulated in *Alternanthera* and *Populus* plants during flood response (Li *et al*., 2017). Another well-studied miRNA, miR156 have been identified for their potential role in submergence and hypoxia by targeting *SQUAMOSA PROMOTER BINDING PROTEIN-LIKE* (*SBP* or *SPL*) genes in lotus and *Arabidopsis* (Moldovan *et al*., 2010, Jin *et al*., 2017, Franke *et al*., 2018). In maize root, miR172 expression was drastically reduced during long-term waterlogging leading to accumulation of its target *AP2/ERF* mRNAs and promotes crown root development (Zhai *et al*., 2013). Comparative analysis revealed that miR775 were suppressed by carbon (C), nitrogen (N), Sulphur (S) and in inorganic phosphate (Pi) deficiency (Liang *et al*., 2015; Kumar *et al*; 2017). Recently, it has been shown that miR775 regulates organ size and reduces cell wall pectin level by negatively regulating their target *GALACTOSYLTRANSFERASE* (*GALT9*) in *Arabidopsis* (Zhang *et al*; 2021). However, their role in submergence is not reported yet.

In the present study, we have explored the role of miR775-*GALT9* module in submergence stress in *Arabidopsis*. We have studied the growth of miR775-Oe1, *galt9*, and *MIM775* lines in submergence recovery to decipher the potential downstream genes. Our study shows the potential involvement of *SAG12*, *SAG29*, *ORE1*, and *RBOHD* genes in imparting submergence tolerance.

## Results

### Expression pattern in co-infiltration experiment and mutant/transgenics, and degradome study showed negative regulation of *GALT9* by miR775

Our degradome analysis identified *GALT9* as a validated target. Cumulative scores for a perfect cleavage site (10^th^) was obtained during degradome analysis through Cleaveland suggesting *GALT9* as a strong target of miR775 in *Arabidopsis*. To further validate the transcriptional cleavage of *GALT9* by miR775, 4-week old *Nicotiana benthamiana* (tobacco) leaves were coinfiltrated with construct *35S:MIR775A* and *35S:GALT9:GFP* (**Figure S1 a**). miR775 coinfiltrated in tobacco leaves will cleave the *GALT9* which will subsequently result in reduced expression of *GFP*. To further estimate the *GFP* level in the tobacco leaves, the tobacco leaves co-infiltrated with construct *35S:MIR775A* and *35S:GALT9:GFP*), we quantified the level of *GFP* by quantitative RT-PCR (qRT-PCR) (**Figure S1 b**). Our result shows that *GFP* expression was significantly downregulated in tobacco leaves co-infiltrated with both the construct (*35S:MIR775A* and *35S:GALT9:GFP*) in comparison to tobacco leaf co-infiltrated with only by *35S:GALT9:GFP*. Reduced expression of *GFP* in tobacco leaf containing both the construct (*35S;MIR775A* and *35S:GALT9:GFP*) confirms the post-transcriptional cleavage of *GALT9* by miR775 (**Figure S1 b**). The coding sequence of *GALT9* consists of 1038 bp including seven exons which are interrupted by 6 introns while GALT9 proteins consist of 345 amino acid residues. GALT9 protein contains terminal glycosyl residue which transfers to its substrate with help of UDP-galactose (Qu *et al*., 2008, Gille *et al*., 2013, Qin *et al*., 2013). *GALT9* shows the closest homology with *Gossypium hirsutum* (cotton) *GhGALT1* and belongs to the Carbohydrate Active enZyme GlycosylTransferases (*CAZy GT*) family (Qin et al., 2013). Total 31 members of the *CAZy GT*-family are present in *Arabidopsis* and out of 31, 20 members have beta (1-3) GTs motif, similar to mammalian systems (Qu *et al*., 2008). In our Standard Nucleotide Basic Local Alignment Search Tool (BLAST) (**Figure S3**), *AT1G53290* showed homology across the Brassicaceae family including *Arabidopsis thaliana, Arabidopsis lyrata subsp. lyrata, Camelina sativa, Capsella rubella, Eutrema salsugineum, Brassica rapa, Brassica napus, Brassica oleracea var. oleracea, Raphanus sativus, Brassica oleracea, Arabis alpina, Brassica rapa subsp. pekinensis, Eucalyptus grandis, Prosopis alba, Camellia sinensis, and Erythranthe guttata* and named as *probable beta-1,3-galactosyltransferase*.

To further confirm the miR775 mediated downregulation of *GALT9* in *Arabidopsis*, we have generated the overexpression construct of *MIR775A* under 35S promoter (*35S:MIR775A*). We checked the expression of miR775 in the T1 plants after selecting the positive from the Hygromycin B supplemented media. We checked the expression of miR775 in the T1 plants selected on Hygromycin B supplemented media. Two lines showing the maximum upregulation of miR775 were miR775-Oe1(14.19-fold) and miR775-Oe2(5.46-fold) (**Figure 1 a**). Both the lines were selected for the further analysis in T3 generation. PCR-based genotyping, the selection of Hygromycin B, and genetic segregation were used to obtain homozygous T3 lines. We have analyzed the expression of *GALT9* in miR775-Oe plants and found the reduced expression in miR775-Oe1(0.52) and miR775-Oe2(0.37) (**Figure 1 b**). Reduction in the expression of *GALT9* in the miR775-Oe line confirms a potential post-transcriptional regulation of *GALT9* by miR775. To further characterize the role of miR775 in *Arabidopsis* development we have used the target mimicry approach (Todesco *et al*., 2010). We have generated a target mimic of miR775 (*MIM775*) transgenic lines which overexpression *GALT9*. The expression of *GALT9* was studied in *MIM775* lines. We have selected the two lines exhibiting the maximum upregulation of *GALT9 MIM775-1* (3.98-fold) and *MIM775-2* (2.55-fold) (**Figure 1 c**). The abundance of *GALT9* transcript in these plants was considerably higher as compared to Col-0, indicating that reduction of miR775 activity leading to the increased expression of *GALT9* (**Figure 1 c**).

**Figure 1.**
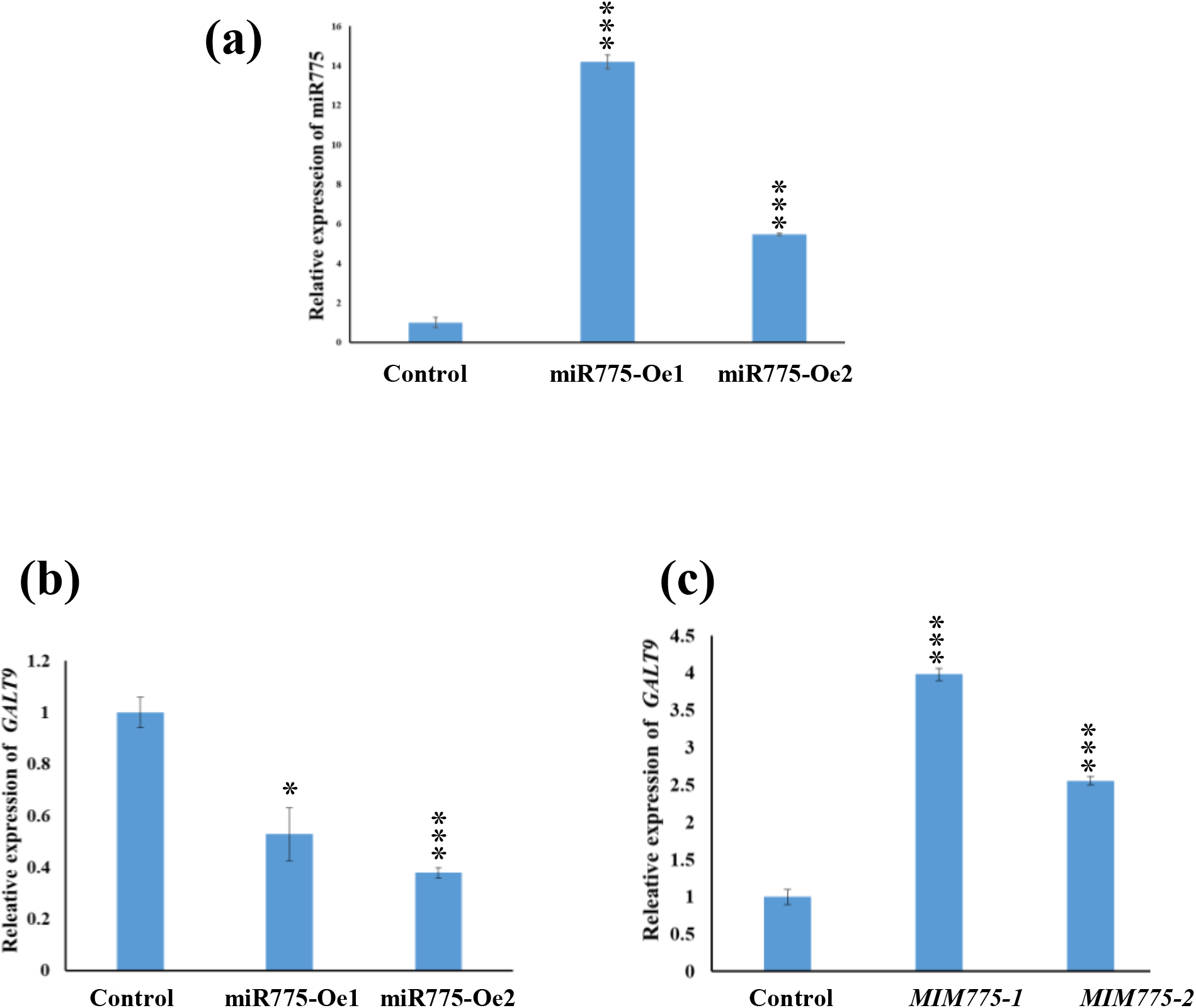
Relative expression of miRNA and their target *GALT9* (a) Relative quantification of miR775 expression in miR775-Oe1 and miR775-Oe2; (b) Relative quantification of *GALT9* expression in miR775-Oe1 and miR775-Oe2; (c) Relative quantification of *GALT9* expression in *MIM775-1* and *MIM775-2*. Error bars indicate standard error (±SE) of three independent experiments. Asterisks indicate significant statistical differences *** P ≤ 0.001, ** P ≤ 0.01, * P ≤ 0.05 using t-test.

### GALT9 protein localizes in Golgi apparatus at subcellular level

To explore subcellular localization of GALT9 protein, we used an *Arabidopsis* Subcellular Localization Prediction Server (AtSubP; http://bioinfo3.noble.org/AtSubP/index.php). For the prediction of GALT9 subcellular localization, we selected amino acid composition-based Support Vector Machine (SVM) and found that GALT9 is a Golgi apparatus protein. To validate the above hypothesis we generated *p35S:GALT9:GFP* construct and co-infiltrated it along with Golgi apparatus marker for transient subcellular localization in *Nicotiana benthamiana* leaves. Differential interference contrast (DIC) microscopy showed that GFP signals were detected primarily in the Golgi apparatus confirming the GALT9 protein localization (**Figure S2**).

### *MIR775A* and target *GALT9* showed complementary expression pattern predominantly in shoot and root tissues

To investigate the expression pattern of *MIR775A* and its target *GALT9* in different developmental stages, we performed the histochemical GUS assay of 5 days after germination (dag) old seedlings and 35 days old plants of *pMIR775A: GUS* and *pGALT9:GUS*. In 5 dag seedlings, the expression of *pMIR775A* was found in the primary root (PR), shoot, hypocotyl, cotyledons, leaves, root shoot junction (RSJ), stomata, and trichomes (**Figure 2 a-f**). The *pMIR775A: GUS* was expressed in young rosette leaves, old rosette leaves, cauline leaves, flowers, and silique in 35 dag plants (**Figure 2 m-p**).

**Figure 2.**
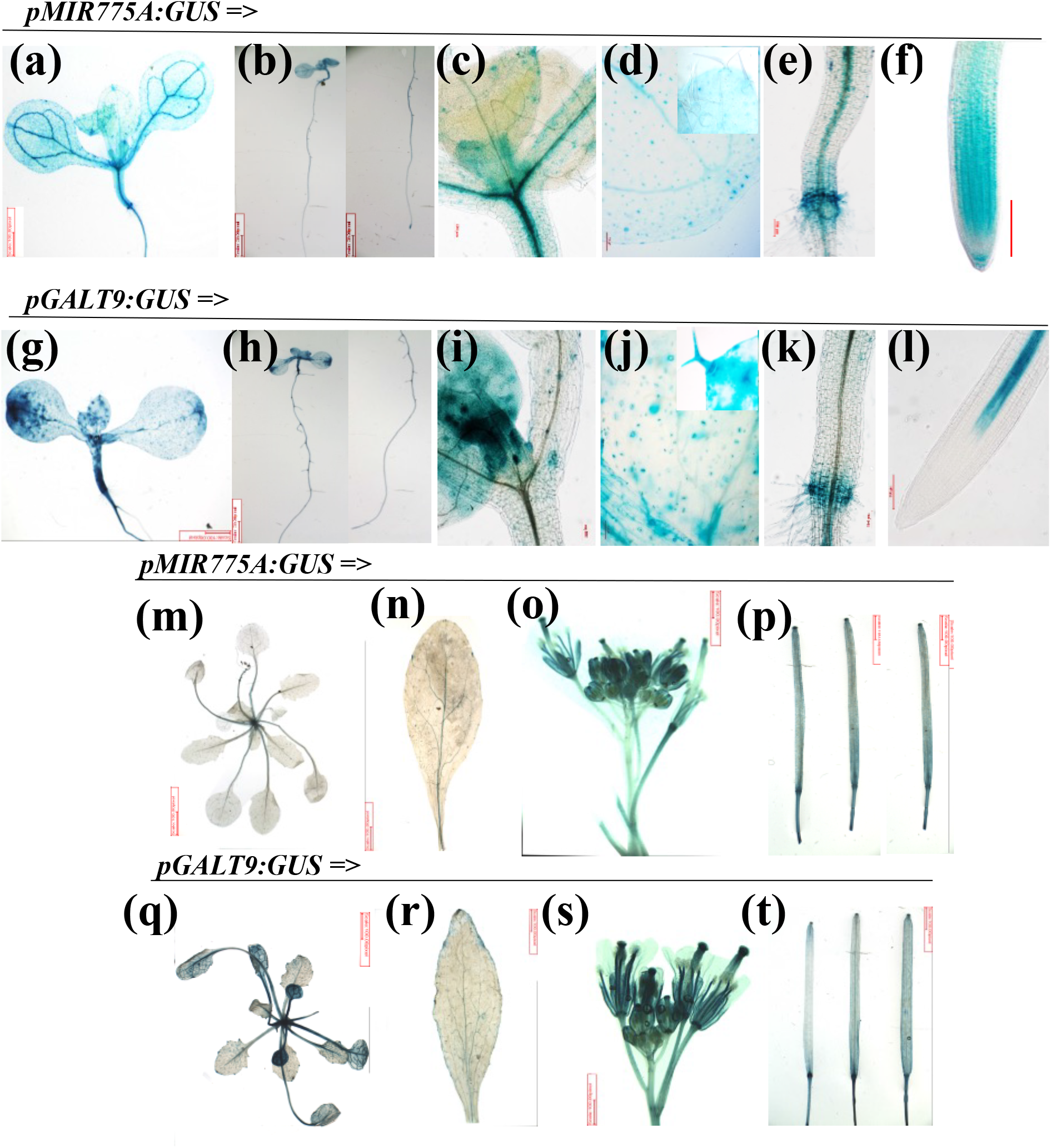
Tissue specific expression pattern of *MIR775A* and its target *GALT9* in seedlings at different time points. (a-f) GUS expression of *MIR775A* in seedlings of 5 days after germination (dag); representation of expressions in different tissues are as following (a) shoot; (b) whole plant & root; (c) shoot apex; (d) leaf; (e) root & shoot junction; (f) root. (g-l) showing GUS expression of target *GALT9*; (g) shoot; (h) whole plant & root; (i) shoot apex; (j) leaf; (k) root & shoot junction; (l) root. (m-p) GUS expression of *MIR775A* in 35 dag of plant; (m) rosette leaf; (n) cauline leaf; (o) flower; (p) silique. (q-t) showing GUS expression of target *GALT9* in 35 dag; (q) rosette leaf; (r) cauline leaf; (s) flower; (t) silique. Scale bar = 100 μm.

The expression of *GALT9* was observed in PR, shoot, root-shoot junction, hypocotyl, leaves, stomata, trichomes in 5 dag seedlings (**Figure 2 g-l**). In addition to this, *pGALT9:GUS* showed expression in rosette leaves, cauline leaves, flowers, and siliques in 35 days old plants (**Figure 2 q-t**). The highest expression of *pGALT9* was observed in leaves and flowers. The GUS expression of *pMIR775A: GUS* and *pGALT9:GUS* were complementary in root-shoot junction, hypocotyl, primary root, stomata, trichome, and rosette leaves.

We performed relative expression analysis of 5 dag seedlings compared with 5 dag shoot and 5 roots (**Figure 3 a**). Interestingly, the results exhibited a higher expression of miR775 in 5 dag shoot as compared to 5 dag root (**Figure 3 a**). qRT-PCR data illustrates the complementary expression of *MIR775A* and *GALT9* in cauline leaves, floral bud, and flower (**Figure 3 b**). The availability of mature miRNA depends on many factors like the activity of miRNA biosynthesis pathway genes, miRNA processing machinery, etc., therefore, we also validated the accumulation of mature miRNA in different tissues of 35 days old col-0 plants by stem-loop qRT-PCR. The mature miR775 was significantly expressed in a young rosette, young cauline, floral bud, and flowers (**Figure 3 b**). These results suggest a dynamic spatial expression pattern of miR775 and its target in *Arabidopsis*.

**Figure 3.**
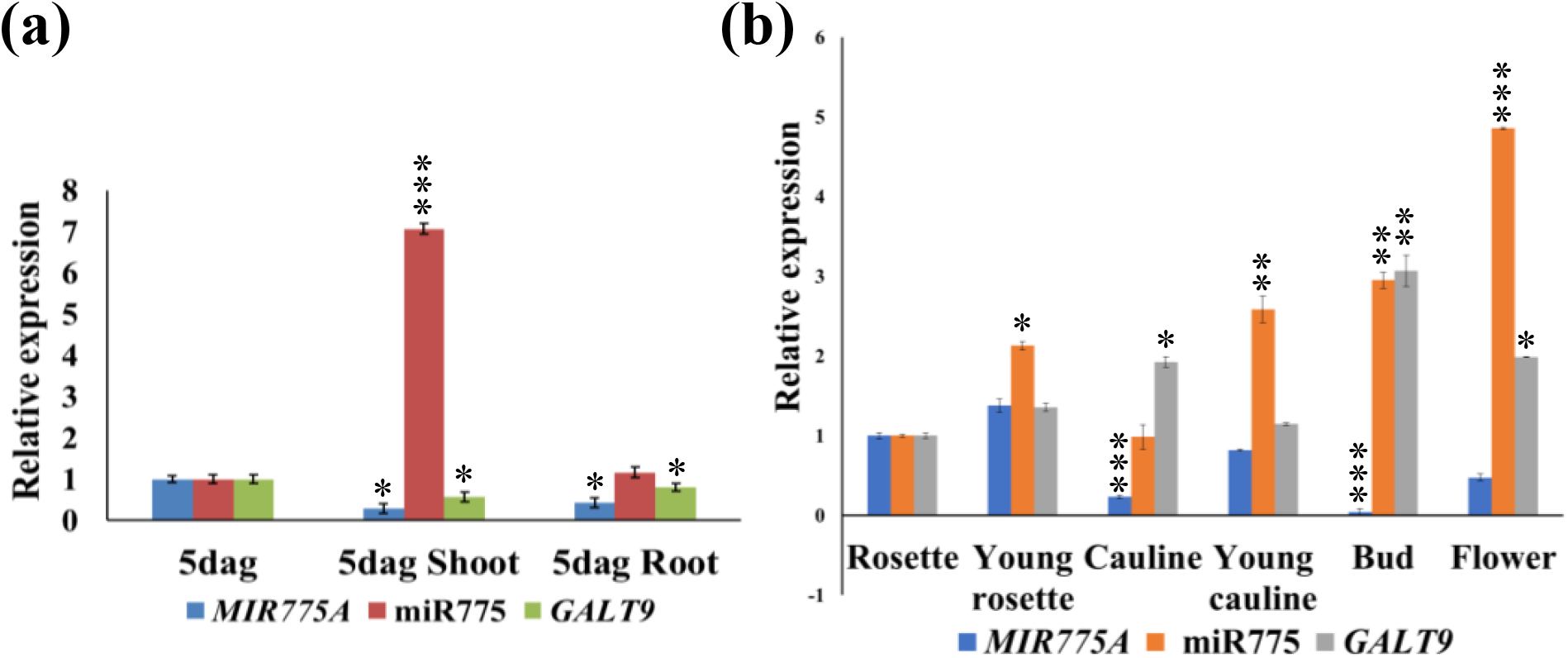
Relative expression of *MIR775A*, miR775 and *GALT9* in various tissues of (a) 5 dag seedlings and (b) 35 dag of plant. Bud (floral bud). Error bars indicate standard error (±SE) of three independent experiments. Asterisks indicate significant statistical differences *** P ≤ 0.001, ** P ≤ 0.01, * P ≤ 0.05 using t-test.

### Submergence stress affects the expression of miR775 and *GALT9* in *Arabidopsis* seedlings

miR775 was reported to be highly upregulated in response to the hypoxic conditions in different microarray and expression studies, in stresses such as flooding and high altitude in *Arabidopsis* (Moldovan *et al*., 2010, Liu *et al*., 2012, Jin *et al*., 2017, Tripathi *et al*., 2019). To analyze the effect on miR775 and its target gene *GALT9* abundance in *Arabidopsis* seedlings during submergence we analyzed the tissue-specific expression of *MIR775* and its target *GALT9* during submergence stress using *pMIR775A: GUS* and *pGALT9:GUS* transgenic plants (**Figure 4 a-j**). Under normoxic (normal level of oxygen) conditions, *pMIR775A: GUS* and *pGALT9:GUS* seedlings showed a basal level of expression in untreated seedlings (0 hr). At 4 hr after submergence *Arabidopsis* seedlings resulted in an initial level of mild elevation of *pMIR775A: GUS* expression however, at 24 hr after submergence the *pMIR775A: GUS* expression was upregulated in shoot tissue (**Figure 4 a-e**). However, at 24 hr after submergence, the *pGALT9:GUS* expression was decreased significantly in shoot tissue (**Figure 4 j**). To validate the submergence induced expression of *MIR775*, we treated 7 dag col-0 seedlings in complete submergence and analyzed the expression of miR775 by stem-loop qRT-PCR and *GALT9* transcripts by qRT-PCR at four different time points (4 hr, 8hr, 12hr, and 24hr) after complete submergence (**Figure 4 k**). Similar to the GUS analysis the expression of miR775 was upregulated by 5 folds at 24 hr however, the expression of its target *GALT9* was downregulated to 0.38 folds at 24 hr of submergence. This result highlights the potential role of miR775 and its target *GALT9* during submergence stress in *Arabidopsis*.

**Figure 4.**
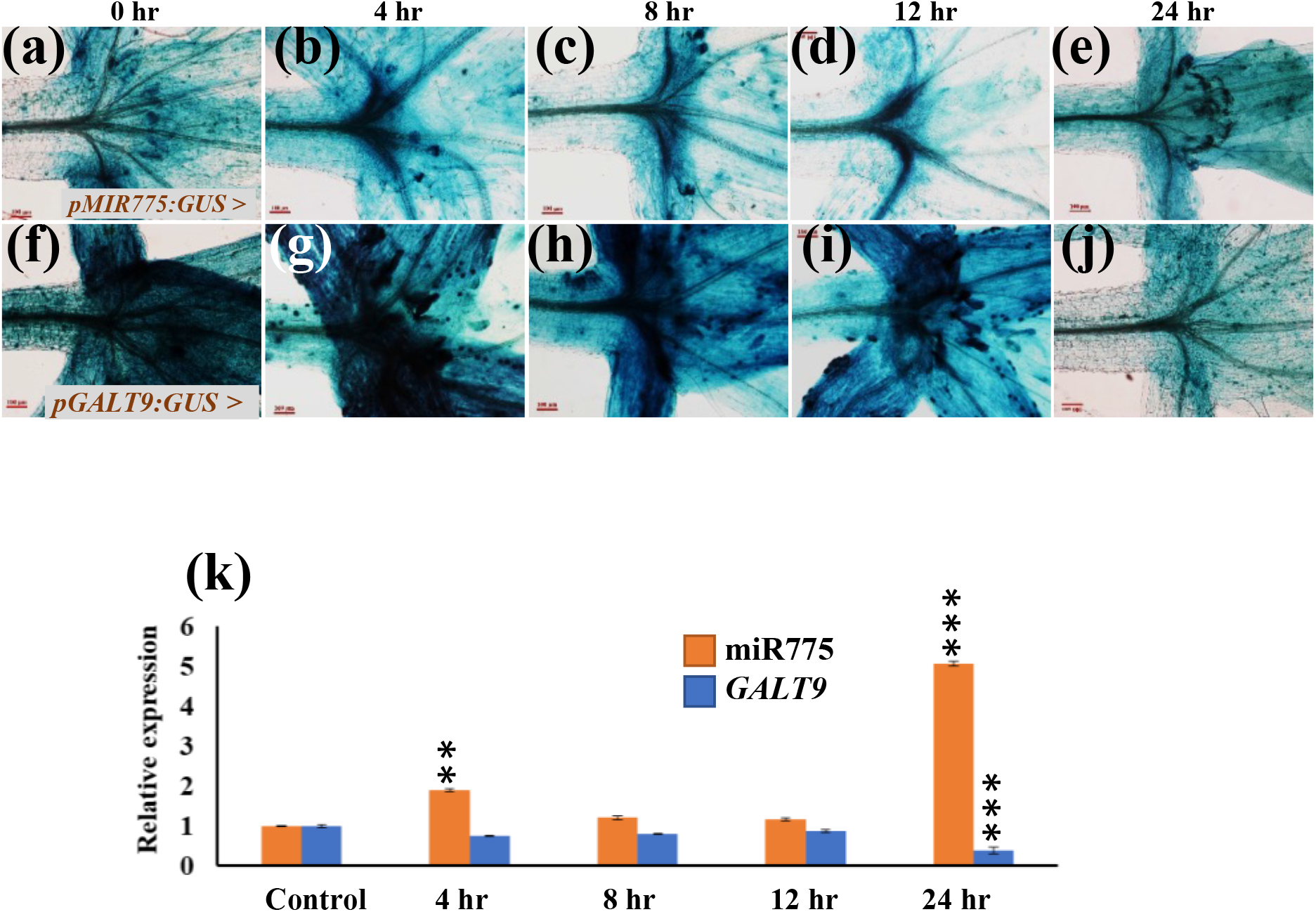
Expression pattern of *MIR775A* and target *GALT9* during submergence stress in *Arabidopsis* seedlings at different time points. (a-e) Tissue specific expression of*pMIR775A:GUS*; (a) control (0 hr); (b) 4 hr; (c) 8 hr; (d) 12 hr; (e) 24 hr. (f-j) Tissue specific expression of target*pGALT9:GUS*; (f) control (0 hr); (g) 4 hr; (H) 8 hr; (i) 12 hr; (j) 24 hr. (k) Relative expression level of miR775 and *GALT9* during submergence stress at different time points. Error bars indicate standard error (±SE) of three independent experiments. Asterisks indicate significant statistical differences*** P ≤ 0.001, ** P ≤ 0.01, * P ≤ 0.05 using t-test. n=10. Scale bar = 100 μm.

### Submergence stress altered the ROS levels in miR775-Oe, *MIM775*, and *galt9* plants

Previously, we have shown that the expression of miR775 and its target *GALT9* was dynamically affected in submergence stress. To further understand the molecular function of miR775 and its target during submergence stress, we further characterized the role of the miR775-*GALT9* module during submergence stress in *Arabidopsis*. Complete submergence leads to the overproduction of different reactive oxygen species (ROS) due to the oxygen-deficient conditions (Bailey-Serres and Voesenek, 2008). Hydrogen peroxide (H_2_O_2_) is one of the major ROS which accumulates during various abiotic and biotic stresses (Liu *et al*., 2010). Increased levels of ROS lead to the impairment in proteins, nucleic acids, lipids, and membrane structures, which eventually triggers cell death (Ghosh *et al*., 2018). ROS level in plants can be estimated through 3,3′-diaminobenzidine (DAB) staining, which is oxidized in the presence of H_2_O_2_ with the help of heme-containing proteins like peroxidases which generates dark brown color (Svistunenko, 2005). Here, we have estimated the ROS levels in different miR775 transgenic lines (miR775-Oe and *MIM775*) and *galt9* mutant seedlings through DAB staining. Among the different transgenic lines, the H_2_O_2_ accumulation was highest in *MIM775-1* and *MIM775-2* (**Figure 6 a-f**). The level of H_2_O_2_ was decreased in miR775-Oe1, miR775-Oe2, and *galt9* as compared to Col-0 (**Figure 6 a-f**). Further, we have also quantified the accumulation of *RBOHD*, a core hypoxia marker gene after the 5 days of desubmergence and we found the expression of *RBOHD* was increased in *MIM775(2.19)* as compared to miR775-Oe(0.01), *galt9*(0.03) transgenics lines (**Figure 6 g**). Increased RBOHD production in *MIM775* suggests ROS accumulation could be persistent even after 5 days of desubmergence, which leads to impaired growth during submergence due to the higher cell death. miR775-Oe lines promote the plant survival ability during recovery from submergence stress due to less ROS levels. Altered ROS levels in different miR775 transgenic and *galt9* lines suggest the different ability of cells to survive therefore, we were interested to estimate the survival ability in different miR775 transgenic lines. To estimate the survival rate, we have treated 25 dag plants of different transgenic lines (miR775-Oe1, miR775-Oe2, *galt9*, and *MIM775-1*, *MIM775-2*) in complete submergence for 5 days and estimated their survival rate after 5 days of desubmergence. Our result indicates that nearly ~75% of the wild type (Col-0) plant survived during submergence recovery, however, the survivability rate was higher in miR775-Oe lines (~85-90%) and *galt9* (~90 %). The survival rate of *MIM775* was reduced to ~50%. (**Figure 5 m**). These results indicate that miR775 mediated regulation of its target gene *GALT9* is potentially involved in the plant survival after submergence stress.

**Figure 5.**
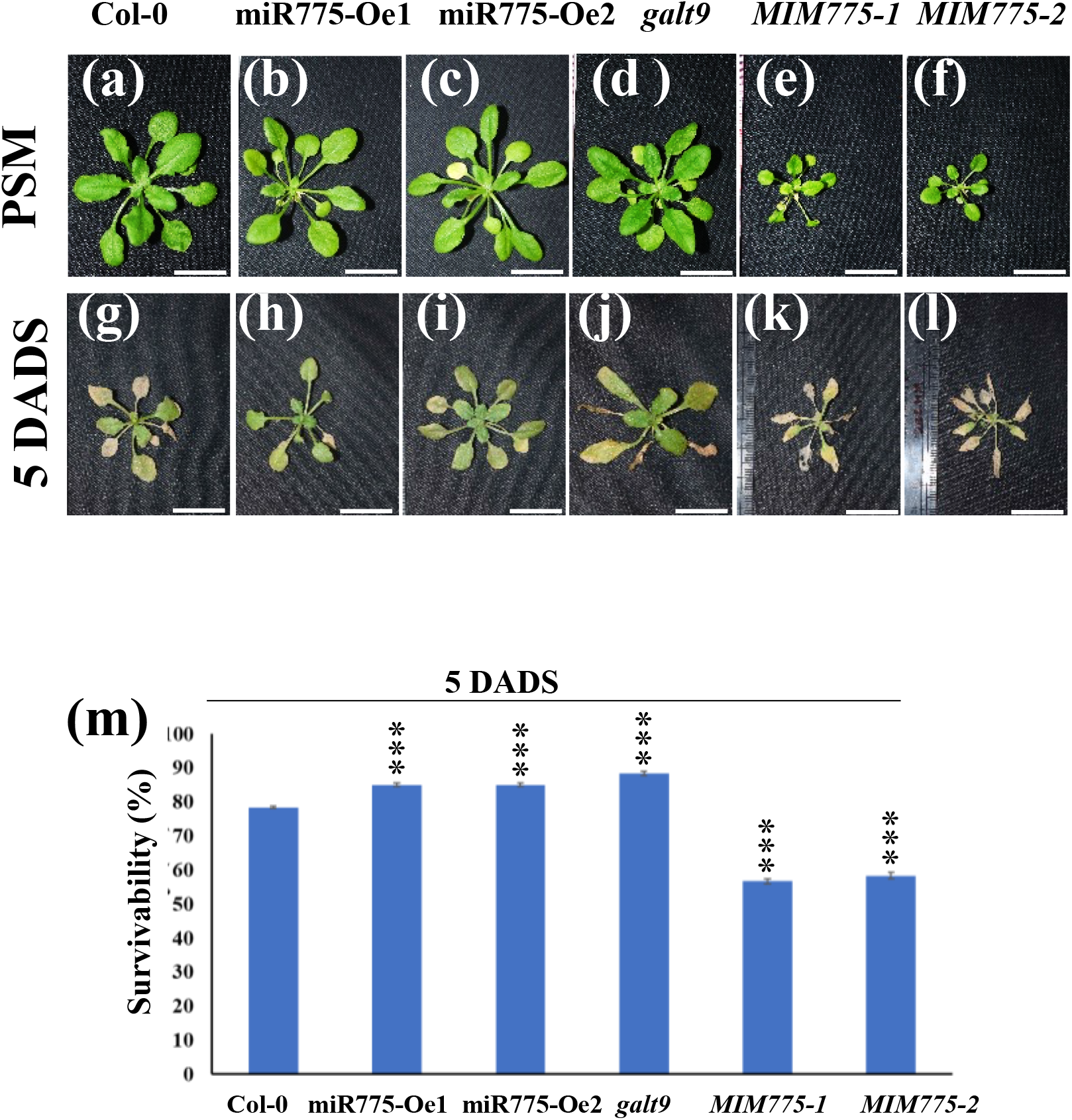
Submergence recovery effect in *Arabidopsis* by miR775 and its target *GALT9*. (a-f) showing phenotypes in different transgenic lines at pre-submergence (PSM) condition of (a) control (Col-0); (b and c) overexpression lines of miR775 (miR775-Oe1 and miR775-Oe2); (d) target mutant *galt9*; (e and f) target mimic lines of miR775 (*MIM775-1* and *MIM775-2*). (g-l) showing phenotypes of different transgenic lines on 5 days after de-submergence (DADS) of (g) Col-0; (h and i) Overexpression lines of miR775 (miR775-Oe1 and miR775-Oe2); (j) target galt9; (k and l) target mimic lines of miR775 (*MIM775-1* and *MIM775-2*). (m) showing Survivability (%) of different transgenic lines of miR775 along with control (Col-0) on 5 DADS.

**Figure 6.**
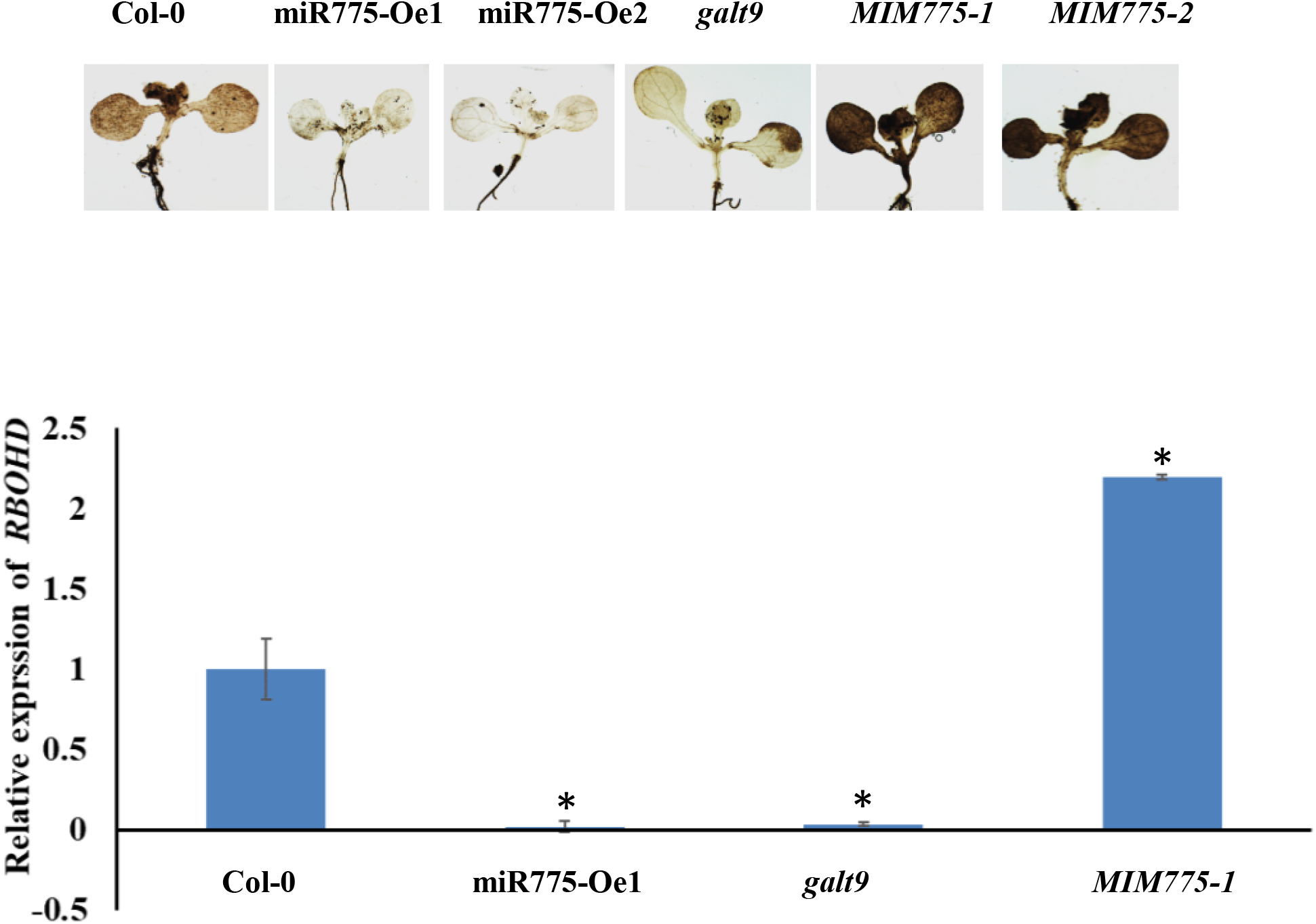
A Controlled ROS production is essential for recovery signaling. (a-g) Representative microscopic images of H_2_O_2_ accumulation (brown color) in (a) Col-0; (b and c) over-expression lines of miR775 (miR775-Oe1 and miR775-Oe2); (d) target *galt9*; (e and f) target mimic lines of miR775 (*MIM775-1* and *MIM775-2*), (g) relative mRNA abundance of *RBOHD* measured by qRT-PCR in Col-0, miR775-Oe1, *galt9* and *MIM775-1* in rosette leaves following 5 days of de-submergence after 5 days of submergence. Error bars indicate standard error (±SE) of three independent experiments. Asterisks indicate significant statistical differences *** P ≤ 0.001, ** P ≤ 0.01, * P ≤ 0.05 using t-test. n=20. Scale bar a-l = 2 cm and n-s =100 μm respectively.

### Expression of senescence-associated genes (SAG) and chlorophyll content were altered post-submergence recovery

Our result showed the poor recovery in *MIM775* lines after submergence in contrast to miR775-Oe and *galt9* lines. The *MIM775* transgenic lines have reduced height and small rosette leaves; however, the miR775-Oe and *galt9* lines were phenotypically similar to the Col-0 under normal growth and developmental conditions. Senescence was promoted in *MIM775* transgenic lines and this *MIM775* also exhibits a high degree of chlorosis after 5 days of post submergence recovery (**Figure 5**). However, the miR775-Oe and *galt9* transgenic lines show reduced senescence and chlorosis as compared to *MIM775* transgenic lines. To further examine the extent of chlorophyll breakdown, we estimated the level of Chlorophyll A, Chlorophyll B, total Chlorophyll, Carotenoids, and Xanthophylls at 5 days after desubmergence (**Figure 7 a-d**). The total chlorophyll level was reduced in *MIM775* transgenic lines in contrast to miR775-Oe and *galt9* transgenic lines. The level of chlorophyll A, B, and xanthophyll was also reduced in *MIM775* lines (**Figure 7 a-d**).

**Figure 7.**
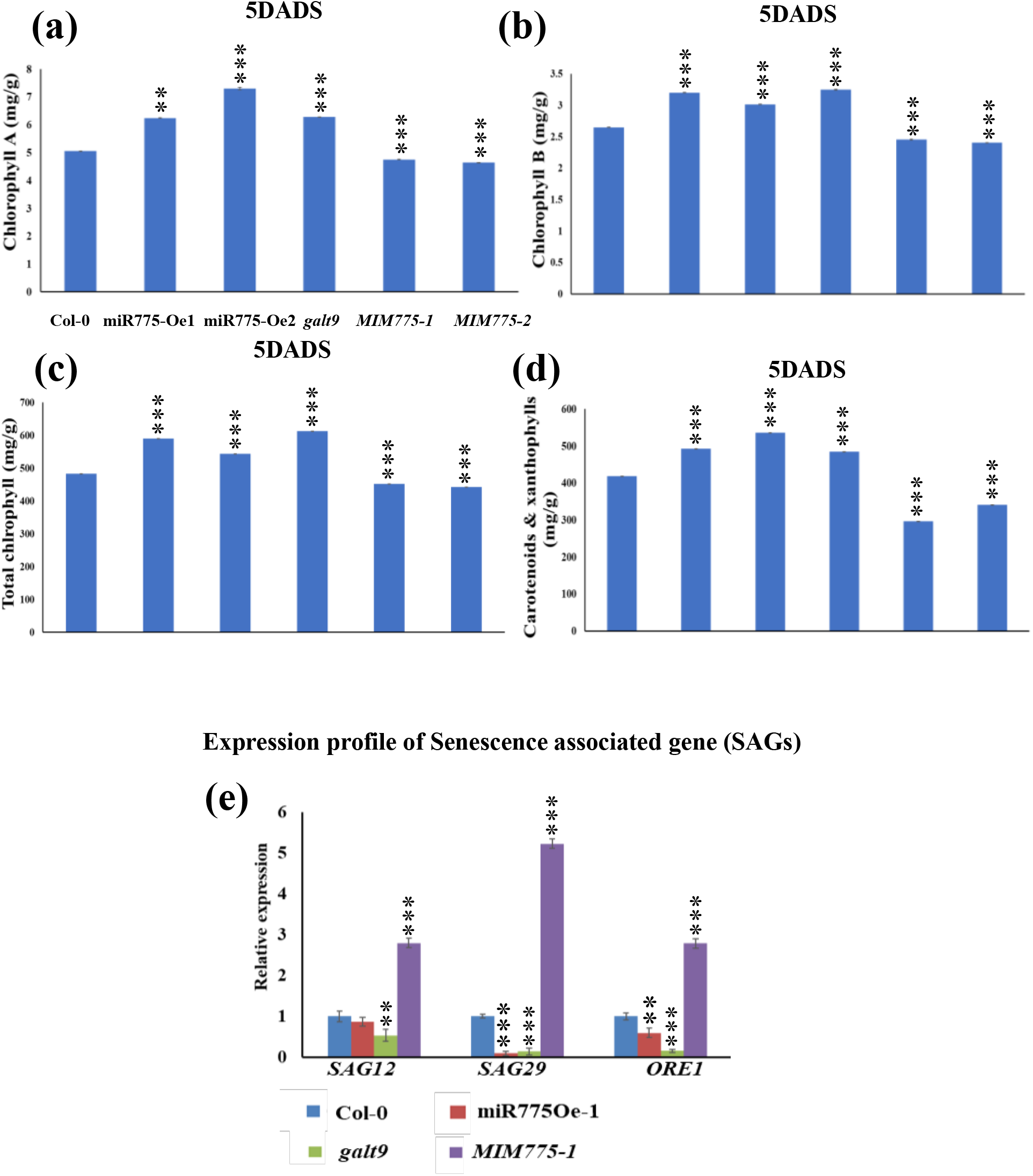
Submergence effect on chlorophyll content and expression of senescence associated genes in miR775 and its target *GALT9*. (a, b, c and d) Estimation of Chlorophyll A, Chlorophyll B, total Chlorophyll, Carotenoids and Xanthophylls respectively in control Col-0, overexpressing line of miR775-Oe1, *galt9* and *MIM775-1* of 5days after de-submergence (DADS); (e) Relative expression levels of senescence associated gene (SAG) *SAG12, SAG29* & *ORE1* respectively in control Col-0, overexpressing line of miR775 (miR775-Oe-1), mutant of target *galt9* (*galt9*) and target mimic line of miR775 (*MIM775-1*) of 5 DADS, Error bars indicate standard error (±SE) of two independent experiment for chlorophyll estimation and three independent experiment for relative expression of qRT-PCR. Asterisks indicate significant statistical differences *** P ≤ 0.001, ** P ≤ 0.01, * P ≤ 0.05 using t-test.

An increase in ROS level and reduction in chlorophyll content after submergence stress is related to precocious senescence. Our results indicate that miR775-Oe and *galt9* plants survive through the submergence stress by decreasing ROS level and reduction in chlorosis, hence, delaying the senescence. On the other hand, *MIM775* plants exhibit early senescence upon submergence stress due to excessive ROS levels and reduced chlorophyll content (**Figure 6 a-f)** and (**Figure 7 a-d**).

A large group of *SENESCENCE ASSOCIATED GENES* (*SAGs*) has been functionally characterized and identified in *Arabidopsis* (Gepstein *et al*., 2003). *SAGs* have been regulated by various environmental signals which include both abiotic and biotic stresses indicating various cross-talks between *SAGs* and stress responses (Lim *et al*., 2007). Genes such as *SAG12, SAG29*, and *ORE1* play an important role in submergence mediated recovery by influencing senescence and chlorophyll breakdown in *Arabidopsis* (Huynh *et al*., 2005, Seo *et al*., 2011a, Yeung *et al*., 2018, Bengoa Luoni *et al*., 2019, Ueda *et al*., 2020). To understand the molecular mechanism underlying leaf senescence in miR775 transgenic and *galt9* lines, we compared the expression level of *SAG12, SAG29*, and *ORE1* during submergence stress.

The expression level of *SAG12, SAG29*, and *ORE1* was downregulated in miR775-Oe1 and *galt9* transgenic lines whereas, upregulated in *MIM775-1* (**Figure 7 e**). These data suggest that miR775 and its target *GALT9* are involved in submergence tolerance through modulating the expression of *SAGs (SAG12, SAG29*, and *ORE1*) in *Arabidopsis*.

## DISCUSSION

Submergence severely affects plant growth and development, survivability, and yield by reducing light intensity, stomatal opening, and gaseous exchange which ultimately affects photosynthesis and respiration. So, timely and efficient recovery from the submergence stress is vital for plant growth and survival. Here we explored miR775-Oe1, *galt9*, and*MIM775* in which differences in their submergence tolerance and recovery were predominantly due to variation in chlorophyll break down and ROS accumulation. Using different transgenics lines such as miR775-Oe1, *galt9*, and *MIM775*, we tried to uncover the molecular, physiological and regulatory components that influence recovery in *Arabidopsis*. It has been previously reported that after desubmergence the reillumination conditions lead to the production of ROS in recovering tissues (Elstner and Osswald, 1994, Smirnoff, 1995). ROS production was different between the miR775-Oe1, *galt9*, and *MIM775* lines, which corresponded to higher *RBOHD* accumulation during recovery in *MIM775*. Even though excessive ROS production is damaging for tissues, controlled ROS production via *RBOHD* might be required for stress signaling during submergence tolerance and recovery. *RBOHD*, a key hypoxia gene, and *ROHBD* mediated ROS burst is crucial for submergence tolerance and recovery (Yeung *et al*., 2018). Balanced ROS production is crucial and needs to be countered by an effective antioxidant mechanism that can control excessive ROS production and associated damage in *Arabidopsis*. However, the recovery signals regulating *RBOHD* remain to be understood. A recent report shows that ABA response in *Arabidopsis* is crucial for submergence tolerance and recovery along with ethylene. Ethylene is a senescence accelerating hormone and promotes the growth and senescence of leaves (Neljubow, 1901, Crocker, 1932, Kim *et al*., 2015, Iqbal *et al*., 2017). It has been shown that ethylene can antagonize ABA action on stomatal opening and closing (Yeung *et al*., 2018). Ethylene accumulation led to a high degree of chlorophyll breakdown and early stomatal reopening in Bay-0 (*Arabidopsis thaliana* accessions). RBOHD-mediated ROS production requires submergence mediated tolerance and recovery which protect against photoinhibition and oxidative damage. ABA and ethylene regulate dehydration, and senescence during submergence recovery by modulating *SAG*s genes and *ORE1*.

Our previous results have indicated the dynamic expression profile of *pMIR775A:GUS* and *pGALT9:GUS* lines during the submergence stress (**Figure 4**). Based on our result (**Figure 4**) and previous reports, we have characterized the miR775-*GALT9* module for its role in submergence stress response. Submergence stress causes oxygen-deprived conditions (hypoxia) (Nishiuchi *et al*., 2012, Ahmed *et al*., 2013, Chen *et al*., 2015, Yeung *et al*., 2018, Loreti and Striker, 2020, Nakamura and Noguchi, 2020). Hypoxic condition is the true consequence of submergence stress. Among the miR775 transgenic lines, miR775-Oe1 and *galt9* lines exhibited the least accumulation of ROS s inferred from our result (**Figure 6 a-f**). However, *the MIM775-1* line displayed a high accumulation of ROS. ROS accumulation indicates the occurrence of cell death during submergence stress. Therefore, our results demonstrate the increased cell death in *MIM775-1* lines in comparison to miR775-Oe1 and *galt9* (**Figure 6 a-f**). Increased cell death in *MIM775-1* during the submergence stress is evident from the seedlings after desubmergences (**Figure 6 a-f**). Before submergence all the miR775 transgenic lines and *galt9* mutants were green and healthy, however, *MIM775* lines were small in size due to the involvement of miR775 in the organ size development through HY5 (**Figure 5 a-f**). Recently miR775-*GALT9* module has been reported to be involved in regulating organ size via HY5. We have also shown the direct binding of HY5 in the promoter of miR775 through yeast one-hybrid assay (**Figure S4**). The miR775-Oe1 and *galt9* transgenics lines were healthy in comparison to Col-0 and *MIM775-1* at 5 days after desubmergences (**Figure 5 a-l**). *MIM775* lines were poorly affected by the submergence stress and showed early senescence during the submergence. The survivability percentage of *MIM775-1* was the least and that of miR775-Oe1 and *galt9* was highest (**Figure 5 m**). These findings strongly indicate the strict regulation of the miR775-*GALT9* module during the submergence stress. Due to submergence stress senescence is induced. Senescence is a well-regulated process of aging in plants. Leaf senescence is directly associated with chlorophyll dissociation. Submergence stress leads to the yellowing of leaves in *MIM775-1* lines therefore we have quantified the chlorophyll content in miR775-Oe1, *galt9*, and *MIM775-1* (**Figure 7 a-d**). Chlorophyll content was least in the *MIM775* lines suggesting elevated senescence in *MIM775* lines (**Figure 7 a-d**). We have further quantified the expression of some key *SAGs* gene *ORE1*, *SAG12*, and *SAG29* through qRT-PCR which is involved in the senescence and leads to chlorosis. Our result illustrated that the expression of these senescence-associated genes such as *ORE1, SAG12, and SAG29* was significantly high in *MIM775* lines in comparison to miR775-Oe1*, galt9*, and Col-0 (**Figure 7 e**). Based on these results it is potentially evident that the miR775*-GALT9* module regulates senescence and stress recovery via *the SAGs* gene which could be either directly or indirectly.

We have also observed the reduction in the size of *the MIM775* shoot in comparison to Col-0, indicating the potential role of miR775-*GALT9* during organ size growth. We have extensively illustrated the role of the miR775-*GALT9* module during submergence induced recovery response. miR775 promotes recovery after submergence by downregulating the expression of *GALT9*. In our study, we have established that *GALT9* overexpression in the *MIM775* line led to the severe senescence in these lines, and due to the enhanced expression of *SAG* genes. Increased expression of *SAGs* and *ORE1* in the *MIM775* lines promotes cell death and chlorosis (**Figure 7 e**). Based on our findings, we propose a model depicting the importance of miR775 in regulating the expression of *GALT9* that regulates submergence recovery (**Figure 8**). miR775-*GALT9* module regulates the recovery from submergence induced senescence by degrading chlorophyll pigments via *SAG*s and *ORE1* either directly or indirectly.

**Figure 8.**
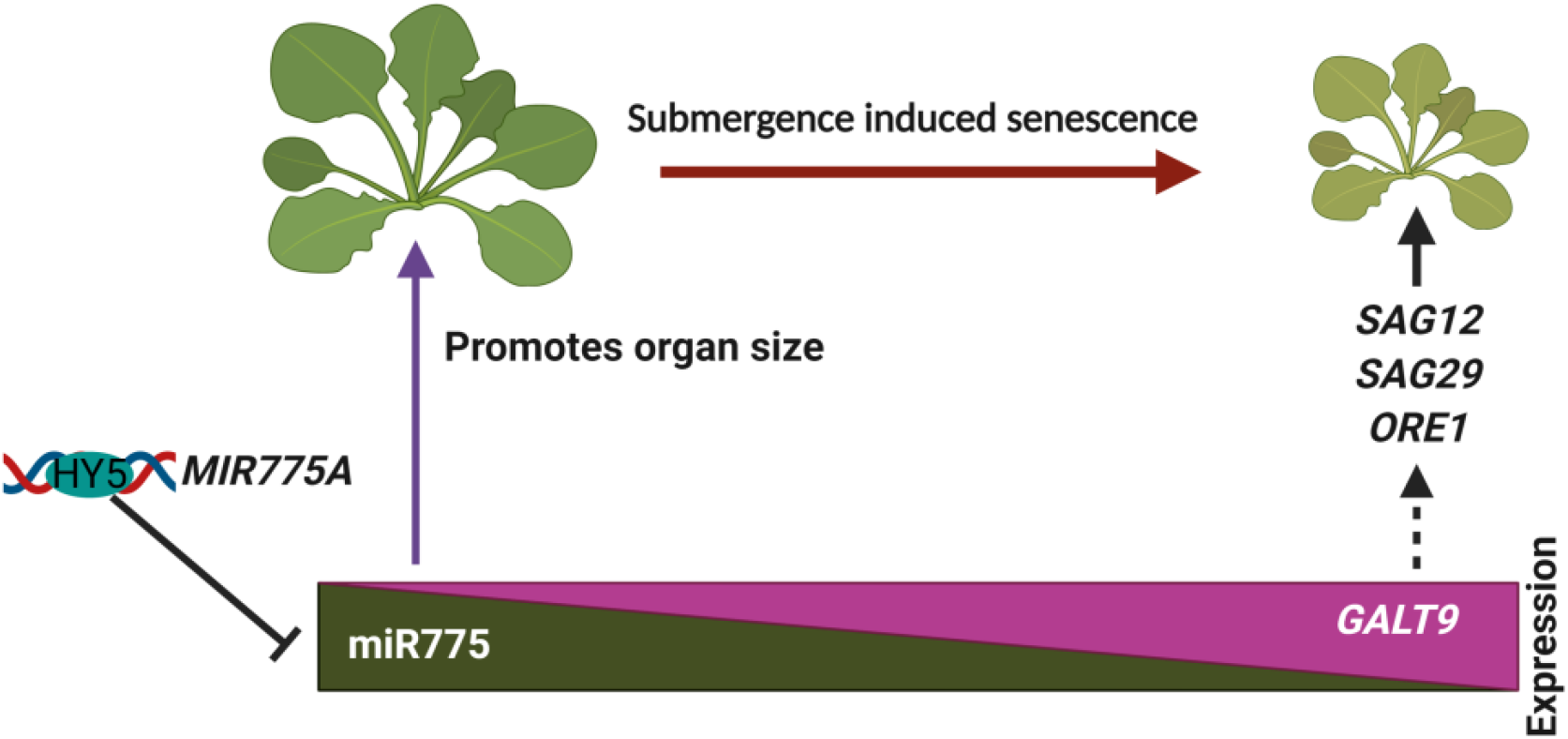
Signaling network mediating post submergence recovery and senescence. *SENESCENCEASSOCIATED GENE* (*SAG12* and *SAG29*) and *NAC* transcription factor *ORE1* accelerates chlorophyll breakdown and senescence.

## Material and methods

### Construction of transgenic lines

For miR775 overexpression, the DNA fragments corresponding to the precursors of 238 bp were cloned, fused to the cauliflower mosaic virus 2X35S promoter gateway cloning vector pMDC32, and transformed into ecotype Columbia (Col-0). For the expression pattern of *pGALT9:GUS*, the promoter sequences of *GALT9*, 1256bp were cloned, fused to the -glucuronidase (GUS) reporter gene like *pGALT9:GUS*, and transformed into Columbia-0 (Col-0) via *Agrobacterium-mediated* transformation. For localization and target validation, the full-length CDS of *GALT9*, 1038bp were cloned in the pCAMBIA1302 vector frame like *p35S:GALT9:GFP* by removing the stop codon. The target mimic line *MIM775* was generated by modifying the *IPS1* gene. *MIM775* target mimic constructs were placed in pGREEN vectors under constitutive CaMV 35S promoter which is resistant against BASTA. Col-0 ecotype was used as control throughout the experiment. Seeds of *galt9* (AT1G53290) SALK_015338 were obtained from ABRC (Ohio State University, Columbus, OH).

### Plant Growth and Submergence Treatment

For all the experiments the *Arabidopsis* seeds were first sterilized by seed wash buffer (70% ethanol and 0.1% (v/v) Triton X-100) and then germinated on half-strength Murashige and Skoog (½MS) medium (HiMedia, Mumbai, India) supplemented with 1% sucrose and 0.8% agar (Murashige and Skoog, 1962). The plants were grown vertically in a controlled environment at 21-22°C, under the 16 h light:8 h dark cycle of white light intensity at 120 μmoles/m2/s for 5days. The above experiments were repeated in triplicates to ensure precision and reproducibility.

For submergence treatment, the disinfected tubs were filled with Milli-Q water overnight before the treatment to maintain temperature equilibrium (21-22°C) was previously described with some modifications (Yeung *et al*., 2018). and submerged (8h after the start of the photoperiod) in ~6 cm water depth in a dark, humidity-controlled condition room. After 5day of submergence, desubmerged plants were replaced under normal growth conditions for 5 days to follow the post submergence recovery. Submergence related experiments were performed at 2:00 PM (8h after the start of the photoperiod).

### Validation of miR775 target using degradome data

We analyzed the target of miR775 through the analysis of *Arabidopsis* degradome PARE data available at Sequence Read Archive (SRA) in NCBI (SRR3143654 – 11 day old seedling, SRR7093799 – leaf sample of stage 5) using tool CleaveLand4 (used to find evidence/s of sliced targets of small RNAs from degradome data) (Addo-Quaye *et al*., 2009). The retrieved SRA datasets, SRR3143654 and SRR7093799, were further converted into fastq and subsequently into fasta file formats using locally installed tools fastq-dump from SRA Toolkit (https://www.ncbi.nlm.nih.gov/books/NBK158900/) and FASTX-Toolkit (http://hannonlab.cshl.edu/fastx_toolkit/), respectively. The tool fastx_trimmer from FASTX-Toolkit was used for trimming the adapter sequences from these datasets. Further, the isoforms of mature miR775 sequences were used as a query against the whole genome cDNA sequences of *Arabidopsis* (https://www.arabidopsis.org/download/index-auto.jsp?dir=%2Fdownload_files%2FSequences%2FAraport11_blastsets). The SRA datasets, mature miR775 sequences and cDNA reference sequences of *Arabidopsis* were used for the analysis of miR775 target through CleaveLand4 pipeline using default settings. The aligned reads of SRA datasets were considered as a target which were being cut at 10^th^ position in miR775.

### *Agrobacterium* infiltration for miRNA target validation

4-week-old *Nicotiana benthamiana* leaves were used for target validation. Oe-miR775 cloned in pSITE-4NB vector and sensitive *35S:GALT9:GFP* constructs were used to transform in *Agrobacterium tumefaciens* GV3101 strain. For infiltrations, the overnight cultures were harvested of individual construct and then suspended in an infiltration buffer (pH 5.7, 0.5% glucose 10 mM MgCl2, 150 μM acetosyringone, and 10 mM MES,) and incubated at room temperature for 6 hrs. For target validation, Nicotiana leaves were infiltrated by 1 ml syringe with the target constructs (sensitive *35S:GALT9:GFP*) alone or in a 1:1 ratio of *35S:MIR775A: 35S:GALT9:GFP* constructs. The plants were kept in a growth chamber maintained at 26 °C (+/−2) and light intensity of 250 μM per m^2^ per s and harvested after 48 hrs for RNA extraction. For target validation, *the HPTII* gene in the vector was used to normalize target abundance in qRT-PCR experiments. For precursor efficiency in qRT-PCR assays, the Ct value of precursor expression was checked to confirm the synthesis of precursor miRNA in the overexpression construct.

### Subcellular localization of GALT9 protein

The coding sequence of the *GALT9* gene (without the stop codon) was cloned into the pCAMBIA1302. The *p35S:GALT9:GFP* construct was then transformed into *Agrobacterium* tumefaciens GV3101 strain. *Agrobacterium* containing the*p35S:GALT9:GFP* or Golgi apparatus marker (GmMan*I*-pBIN2) mCherry constructs were grown to saturation in Luria-Bertani (LB) medium. Cultures were centrifuged and resuspended in 10 mM MgCl2, 10 mM MES, and 150 mM acetosyringone and kept at room temperature for 2 h. The cultures were then diluted to 1 OD600 unit and co-infiltrated into the abaxial side of young tobacco (*Nicotiana benthamiana*) leaf epidermis (four-week-old seedlings grown at 22° C) using a 1 ml syringe without the needle. Transformed leaves were analyzed 72 h after infection of the lower epidermis. Subsequently, fluorescence microscopy was performed on a Nikon 80i as employed to record and process the digital images. At least three independently transformed leaves were analyzed.

### Histochemical detection of GUS assay and microscopy

Histochemical GUS analysis was carried out by putting the samples into appropriate amounts of GUS histochemical GUS buffer (50 mM sodium phosphate, pH 5.7,50 mM EDTA, pH 8.0; 0.1% Triton X-100, 2mM potassium ferrocyanide, 2mM potassium ferricyanide and 1mM 5-Bromo-4-chloro-3-indolyl-б-D-glucuronic acid (X-Gluc) and incubated at 37°C for 14 hrs and 30 mins (mins). Stained samples were washed with a de-staining solution having ethanol: acetone: glycerol (3:1:1 v/v) to remove chlorophyll and then microscopy was performed after putting the samples in chloral hydrate (TCI Chemicals) for 1 hr on a Nikon80i and Olympus SZX16 as employed to record and process the digital images.

### Conditions for the Expression pattern of *MIR775A* and target *GALT9* during submergence stress in *Arabidopsis thaliana s*eedlings at different time points

For submergence treatment, the disinfected tubs were filled with Milli-Q water overnight before the treatment to maintain the temperature equilibrium (21-22°C). The 7 dag seedlings of *pMIR7775A: GUS*, *pGALT9: GUS*, and Col-0 plants containing half MS media in disposable, square plate Petri dishes (120X120X17) mm ht. (company Praveen Scientific Corporation) were used throughout the experiment. The plants were submerged (8h after the start of the photoperiod) at 2:00 PM in ~6 cm water depth in a dark, humidity-controlled condition room. The samples were harvested for GUS analysis at different time points of 4 hrs, 8 hrs, 12 hrs, and 24 hrs. For control plants, the samples were kept in dark for the same period without giving submergence to nullify the effect of the dark with submergence, and then samples were harvested for GUS analysis. For GUS analysis samples were incubated at 37°C for 14 hrs and 30 mins.

### Total RNA Extraction and Quantitative Real-Time qPCR

Gene-specific primers were designed using IDT software Inc. (USA) and custom synthesized from Sigma Aldrich. The total RNA was extracted by using TRIzol (TRI reagent). Purified RNA used for single-stranded cDNA was synthesized from 2.5 μg RNA using oligo(dT) primer by high-capacity cDNA reverse transcription Kit (Thermo Fisher Scientific). The RT reaction consisted of total RNA, 0.8 μl of 100 mM dNTP mix, 2 μl of 10X reaction buffer, 1 μl of random hexamer primer, 1 μl of oligodT, and 1 μl of Revert aid RT enzyme in a final volume of 20 μl. The reaction was carried out at 25°C (10 mins) / 37°C (2 hrs) followed by denaturation at 85°C for 5 mins. For performing qRT-PCR, the cDNAs were diluted to 20 ng with sterile MQ water. For each tissue type, separate PCR amplification reactions were set up for detecting different genes. The qRT-PCR reaction was set up by mixing 5 μl of 2X Power Syber^®^ Green PCR Master Mix (Applied Biosystems, USA), 0.5μl of 10 μM each of forward and reverse primers, 2μl (40 ng) of cDNA and sterile MQ water to adjust the reaction volume to 10μl. qRT-PCR was carried out in Applied Biosystems ViiA 7 Real-Time PCR System with PowerUp^™^ SYBR™ Green Master Mix (Applied Biosystems™ by Fisher Scientific). The relative transcript level was calculated by using the 2-ΔΔCT method which was normalized to *ACTIN2* and *PROTEIN PHOSPHATASE 2A SUBUNIT A3* s (*PP2A3*), as previously described by (Wang *et al*., 2014, Singh *et al*., 2020b).

For stem-loop cDNA synthesis total of 200ng of purified RNA was taken then mix with 0.5 μl 10 mM dNTP and nuclease-free water, incubating the mixture at 65°C for 5 mins and put on ice for 2 mins, brief centrifugation, and add, 4 μl 5× buffer (250 mM Tris-HCl (pH 8.3), 250 mM KCl, 20 mM MgCl2, 50 mM DTT 2 μl 0.1 M DTT) 0.25 μl RiboLock RNase Inhibitor (20 units/μl) 0.25 μl RevertAid H Minus M-MuLV Reverse Transcriptase (200 units/μl), 1 μl Stem-loop primer (1 μm) mix gently and centrifuge to bring the solution to the bottom of the tube and cDNA synthesis was performed by previously described by (Varkonyi-Gasic *et al*., 2007). The cDNA was diluted up to 2 times before performing the real-time PCR. Fold change was calculated using formula FC = 2-ΔΔCt as previously described by (Singh *et al*., 2017, Gautam *et al*., 2020, Singh *et al*., 2020b).

### Histochemical detection of H_2_O_2_

The H_2_O_2_ staining agent, 3,3′diaminobenzidine (DAB) (SRL), was dissolved in H2O by adjusting the pH to 3.8 with KOH. Freshly prepared DAB solution was used to avoid auto-oxidation. The 7 days old seedling were transferred for 5 days for submergence, after 5 days of submergence the seedling was exposed to 1 hour of normal condition and then transferred for treatment and were immersed and infiltrated under vacuum with 1.25 mg ml-1 DAB staining solution for 15 mins and incubated at room temperature for 6 hrs. The Stained seedlings were then bleached out in ethanol: acetic acid: glycerol (3:1:1) (v/v/v) solution for 30 mins and then images were taken by an Olympus SZX16 microscope. The brown color visualization of H_2_O_2_ was due to DAB polymerization.

### Chlorophylls and xanthophylls estimation

Chlorophyll and xanthophyll were extracted from 100 mg of rosettes leave with 96% (v/v) by DMSO dark incubated at 65 °C for 30 mins and cooled to room temperature for 15 mins [149, 150]. Measuring the absorbance at 470, 645, and 663 nm by calibrating to zero with pure DMSO and was measured with a spectrophotometer plate reader (Synergy HT Multi-Detection Microplate Reader; BioTek Instruments). Measurements were done in 2 biological replicates. Chlorophyll A and B concentrations were calculated by the following Arnon‘s equations:

Chlorophyll A (mg/g fresh weight) = [(12.7 X A663) -(2.69 X A645)]X (V/1000 X W) Chlorophyll B (mg/g fresh weight) = [(22.9 X A645) -(4.68 X A663)]X (V/1000 X W) Total Chlorophylls Content= (2008 X A645+802 X A663) X (V/1000 X W) Carotenoids + xanthophylls (mg/g fresh weight) = (1000 X A470-1.90ChlA-63.14ChlB/214) X (V/1000XW)

Where;

V= volume of extract (ml)
W= fresh weight of the sample (g)

### Yeast one-hybrid assays

Yeast one-hybrid assays (Y1Hs) were performed to verify the gene-gene interactions, using the MatchmakerTM Gold Y1H Library Screening System. The full-length CDS of *HY5* was subcloned into the *pGADT7* AD vector and the promoter of *pMIR775A* (~152 bp) was constructed into the public vector according to the ClonExpress II One-Step Cloning Kit. Autoactivation and then interaction analyses were performed.

### Determination of autoactivation concentration of AbA

A healthy colony was picked from the bait strains. The colony was resuspended in SD-Ura broth. The dilution was adjusted to 0.1 0.01 0.001 0.0001 and 10μl of the culture was patched on the following media. Colonies were grown for 2–3 days at 28 °C on SD/-Ura plates.

1. SD/-Ura with AbA (150 ng/ml)
2. SD/-Ura with AbA (250 ng/ml)
3. SD/-Ura with AbA (500 ng/ml)

Vector/Construct details

1. pGADT7 (For Cloning of Prey)
2. pAbAi (For Cloning of bait)
3. pGADT7-Rec-p53/p53-AbAi (positive Control)
4. pGADT7 transformed in Y1H gold cells (negative control).

## ACCESSION NUMBERS

*Arabidopsis* Genome Initiative (AGI) locus identifiers for the genes mentioned in this article are listed as follows:

*MIR775A* (AT1G78206), *GALT9* (AT1G53290), *SAG12* (AT5G45890), *SAG29* (AT5G13170), *ORE1* (AT5G39610), *ROHBD* (AT5G47910), HY*5* (AT5G11260).

## ACKNOWLEDGMENTS

We acknowledge NIPGR for providing necessary research facilities and internal grants. We also thank DBT-eLibrary Consortium (DeLCON) for providing access to e-resources. We acknowledge the Department of Biotechnology (Govt. of India) for providing fellowships.

## AUTHOR CONTRIBUTIONS

VM performed and designed most of the experiments.VM, ArS and NG prepared manuscript draft. VM, ArS, NG, SSD, SY and AK contributed to the manuscript writing and revising the manuscript. AKS has conceptualized the work, interpreted data, and revised the manuscript.

## Supplemental Figure

**Figure S1:**
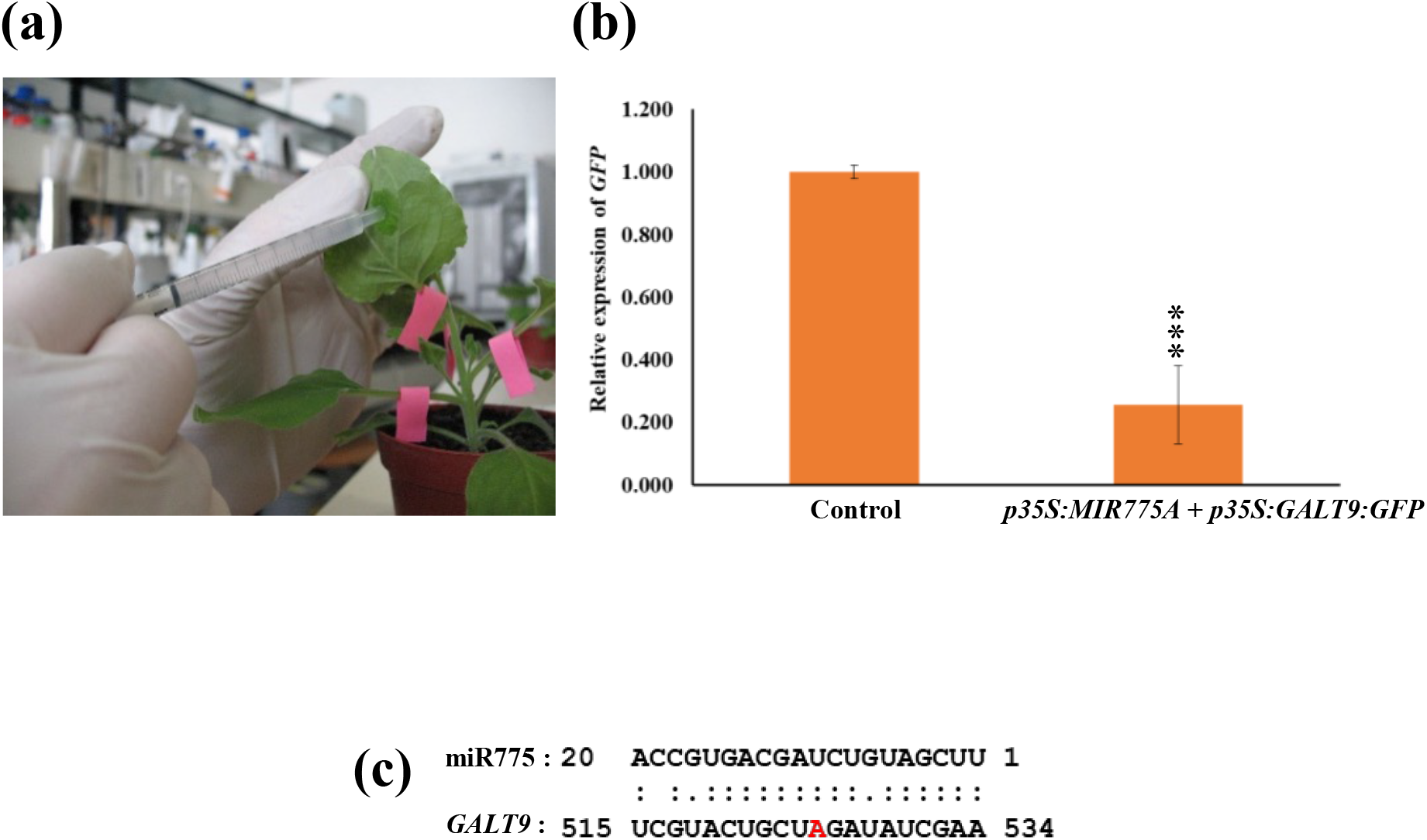
miR775 *c*leaves its predicted target *GALT9* in a transient assay carried out in tobacco (*N. benthamiana*) leaves. (a-f) showing, (a) Co-infiltration of *MIR775A* and *GALT9 (p35S:MIR775A* + *p35S:GALT9:GFP*) in leaf of tobacco plants (~4 weeks old); (b) Relative expression level of *GFP* in control (*p35S:GALT9-GFP*) and *p35S:MIR775A + p35S:GALT9-GFP*; (c) Sequence alignment of the mature miR775 with the target *GALT9*.

**Figure S2:**
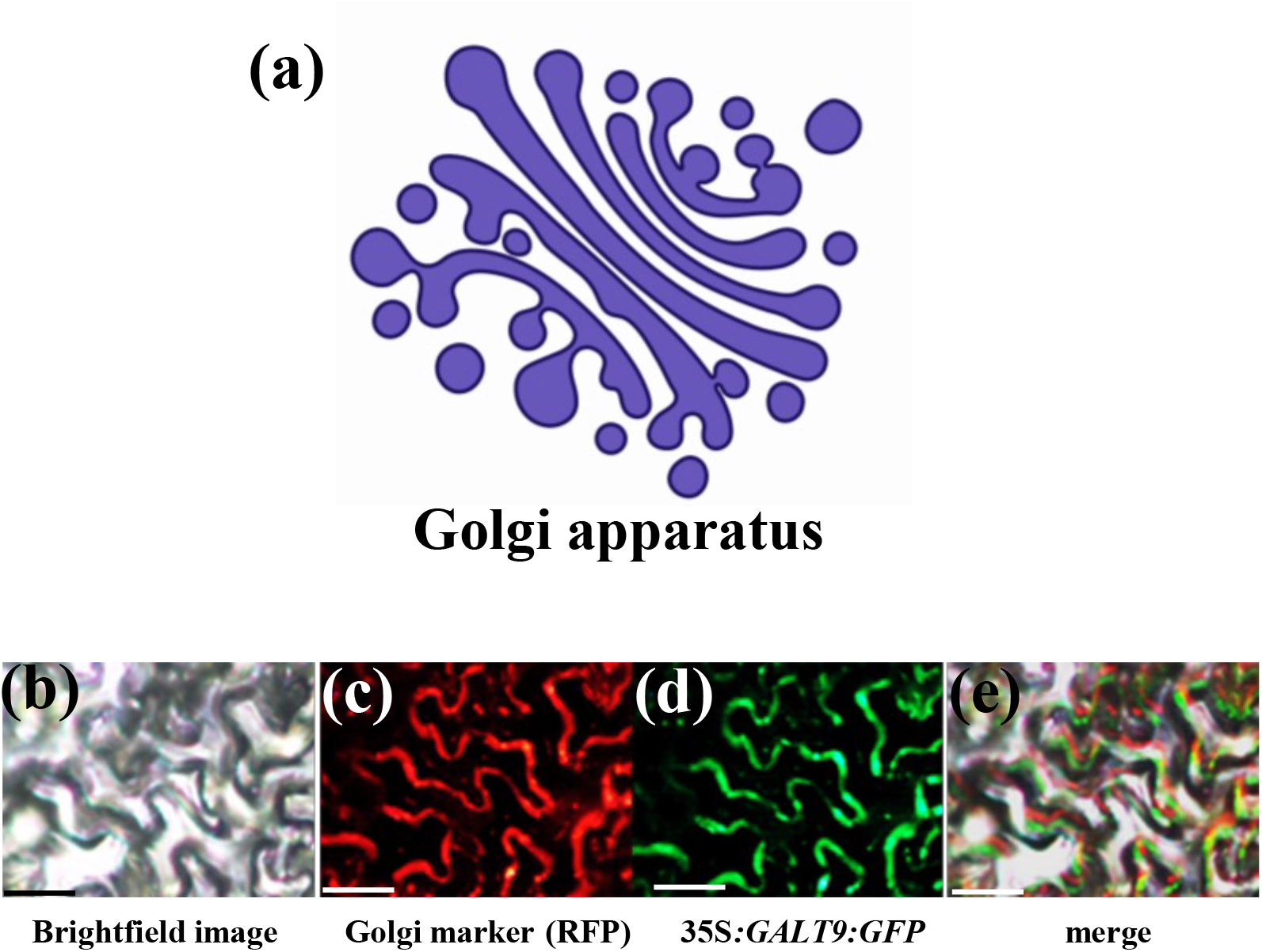
GALT9 localizes in the Golgi apparatus of a cell. (a) The prediction of trans-membrane helices in GALT9 protein (http://bioinfo3.noble.org/AtSubP). (b to e) The fluorescent protein-tagged GALT9 fusion proteins were co-expressed with GmMan*I*-pBIN2 (mCherry Golgi apparatus marker) into the abaxial side of a young in tobacco (*N. benthamiana*) leaf epidermis. The signals were visualized under a fluorescence microscope. (b) Bright field image of leaf epidermal cells; (c) Fluorescence image of *GmManI-pBIN2* (Golgi apparatus marker); (d) Fluorescence image of *35S:GALT9:GFP*; (e) Image was merged with fluorescence image of GmMan*I*-pBIN2 and*p35S:GALT9:GFP*. Scale bar = 50 μm.

**Figure S3:**
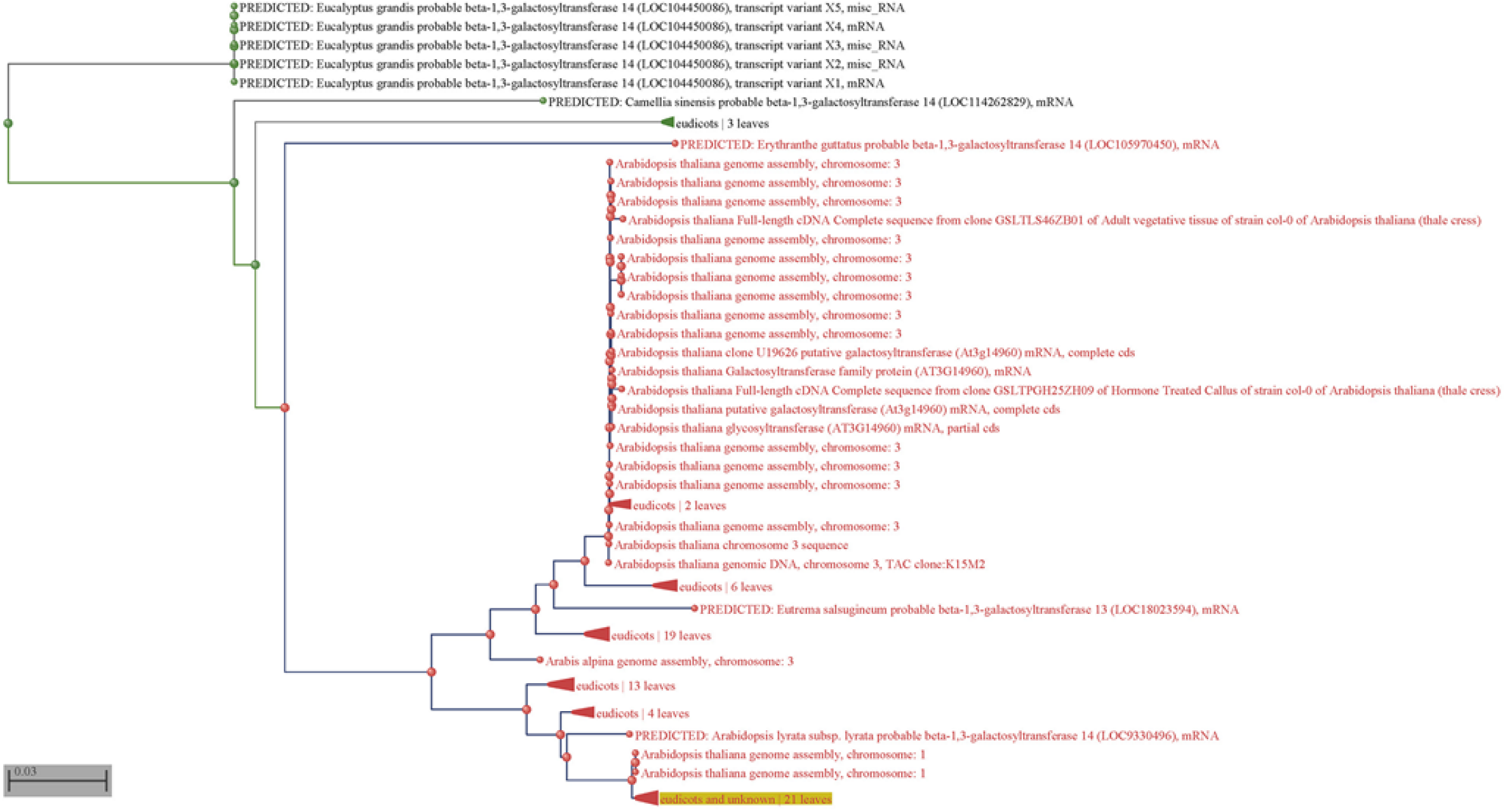
Phylogenetic similarity of At1g53290.

**Figure S4:**
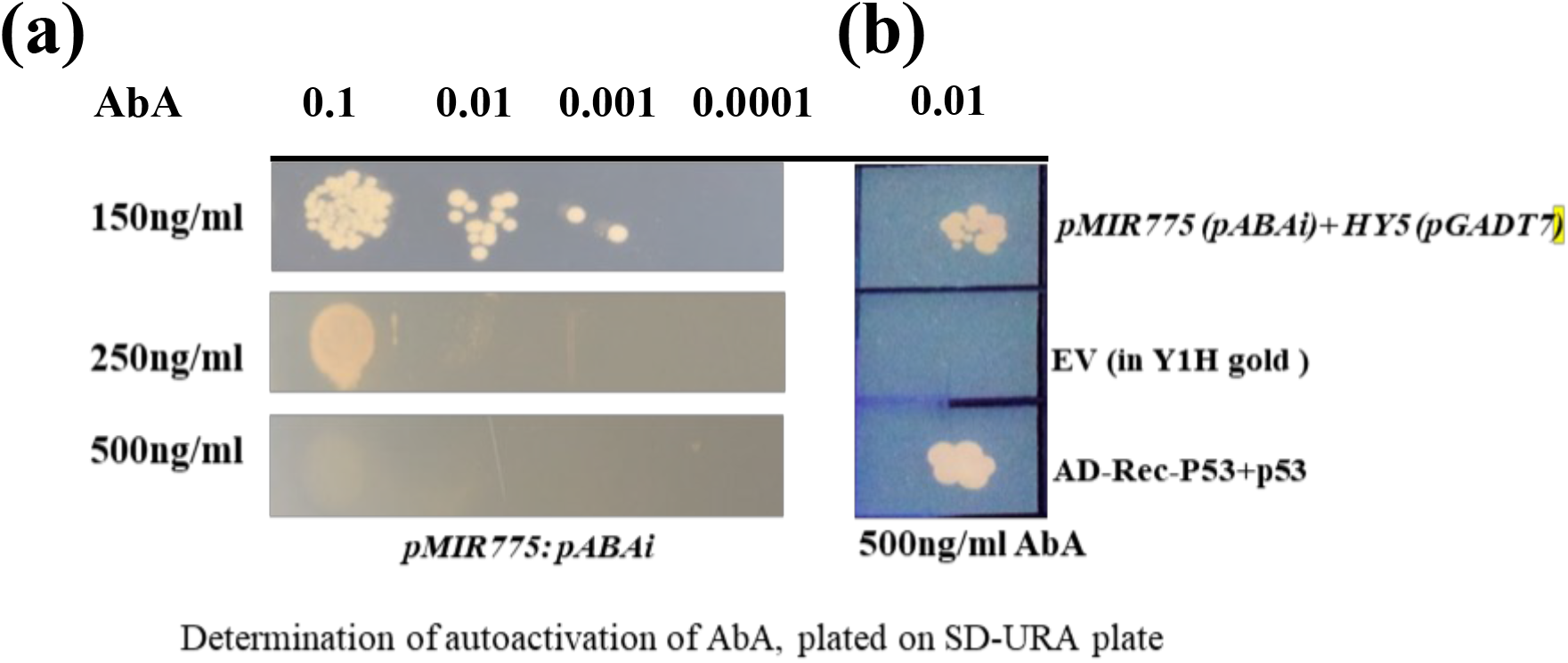
HY5 binds to the promoter of *MIR775A* and positively regulates its expression. (a-c); (a) determination of autoactivation; (b) Represents Yeast one hybrid (Y1H) assay showing the interaction of HY5 with*pMIR775A. HY5* CDS + *pGADT7* transformed into pMIR775A + pAbAi was used to check the interaction. pGADT7-Rec-p53/p53-AbAi was used as a positive control. pGADT7 transformed into a Y1H gold cell used a negative control.

